# SMAD4 and TGFβ are architects of inverse genetic programs during fate-determination of antiviral memory CD8 T cells

**DOI:** 10.1101/2021.12.16.472993

**Authors:** K. Chandiran, J.E. Suarez-Ramirez, Y. Hu, E.R. Jellison, Z. Ugur, J.S. Low, B. McDonald, S.M. Kaech, L.S. Cauley

**Affiliations:** Department of Immunology, UCONN Health, Farmington, CT. USA.; Department of Microbiology and Immunology, Emory Vaccine Center. Emory University, GA. USA; Department of Immunobiology, Yale University School of Medicine, New Haven, CT, USA; NOMIS Center for Immunobiology and Microbial Pathogenesis, The Salk Institute for Biological Studies, La Jolla, CA, USA

**Keywords:** Respiratory infection, cytotoxic T lymphocytes, memory CD8 T cells, homing receptors, transcription factors.

## Abstract

Transforming growth factor β (TGFβ) is a morphogenic protein that augments antiviral immunity by altering the functional properties of pathogen-specific memory CD8 T cells. During infection TGFβ inhibits formation of effector (TEFF) and central memory CD8 T cells (TCM), while encouraging tissue-resident memory CD8 T cells (TRM) to settle in peripheral tissues. SMAD proteins are signaling intermediates that are used by members of the TGF cytokine family to modify gene expression. For this study, RNA-sequencing was used to explore how regulation via SMAD4 alters the transcriptional profile of antiviral CTLs during infection. We show that SMAD4 and TGFβ cooperatively regulate a collection of genes that determine whether specialized populations of pathogen-specific CTLs circulate around the body, or settle in peripheral tissue. The target genes include multiple homing receptors (CD103, KLRG1 and CD62L) and transcription factors (Hobit and EOMES) that support memory formation. While TGFβ uses a canonical SMAD-dependent signaling pathway to induce CD103 expression on TRM cells, an alternative SMAD4-dependent mechanism is required for formation of TEFF and TCM cells in the circulation.

**Graphical abstract:** TGFβ and SMAD4 modulate gene expression in reciprocal directions during differentiation of antiviral CTLs.

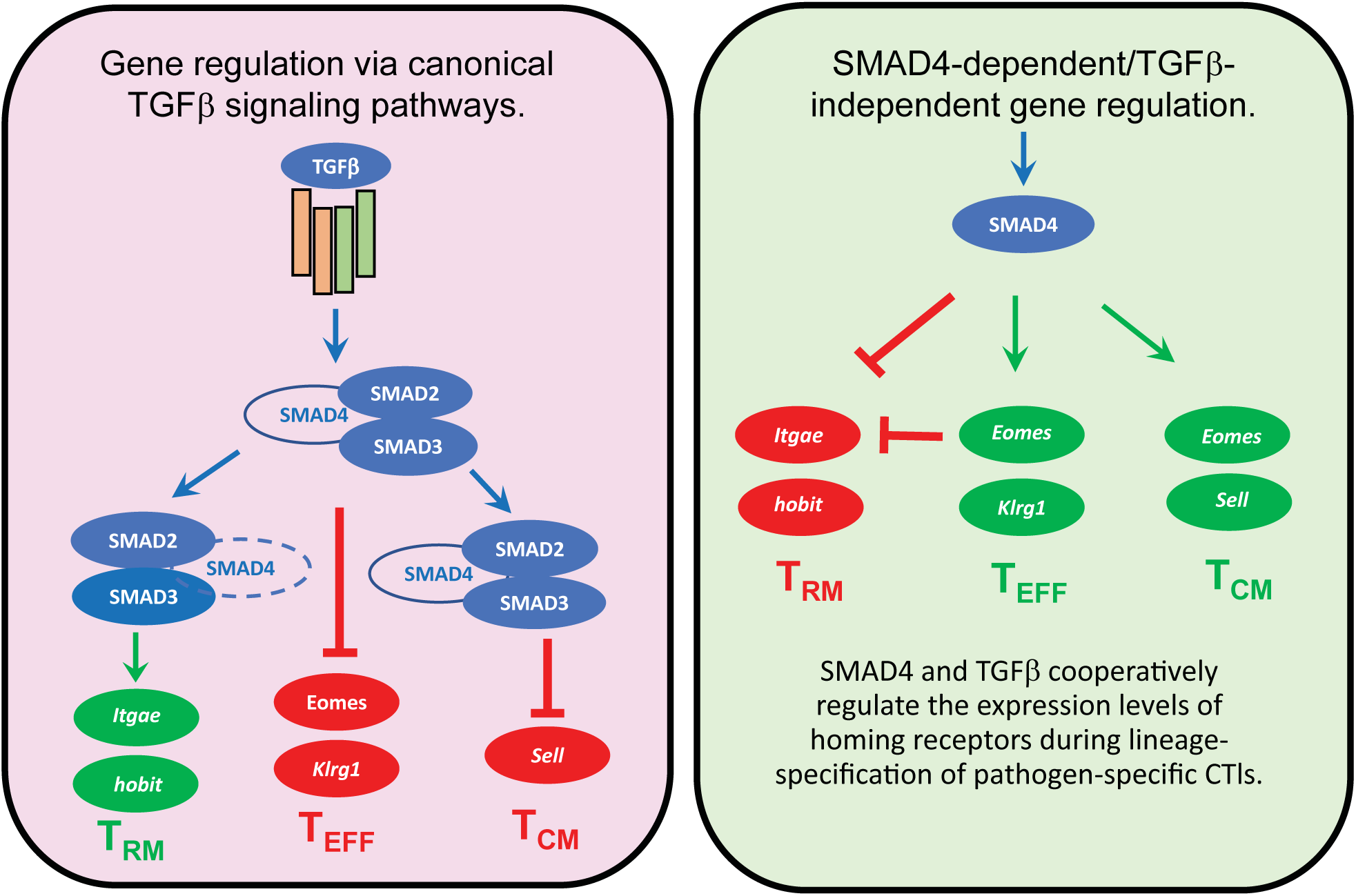

## Introduction

Influenza viruses are highly contagious pathogens that enter the body via the respiratory tract. Cytotoxic T lymphocytes (CTLs) help control these infections by eliminating cells that support the pathogen’s lifecycle. During infection, naïve CD8 T cells differentiate into multiple subsets of effector (TEFF) and memory CD8 T cells that perform specific functions during the immune response. These subsets can be distinguished using surface receptors that are required for effector functions, prolonged survival and localization in different anatomical niches. Although many inflammatory mediators influence the fate decisions of activated CTLs, the mechanisms that coordinate changes homing receptor expression as CTLs respond to during infection are poorly understood. We previously found that a network of signaling pathways, that are controlled by members of the transforming growth family (TGF), alter the functional attributes of pathogen-specific CTLs during memory formation (Cao et al., 2015; Hu et al., 2015b). For the current study, we used additional stains of genetically- modified mice to further define how these important regulatory pathways influence genetic-programming of pathogen-specific CTLs.

To augment immunity, pathogen-specific CTLs must distribute to peripheral and lymphoid tissues. A response to infection is initiated by naïve CD8 T cells that migrate through secondary lymphoid organs (SLO). Circulating lymphocytes enter resting lymph nodes from wide blood vessels known as high endothelial venules (HEV) (Mionnet et al., 2011). During transit through HEVs, CD62L interacts with carbohydrates expressed on the surface of activated endothelial cells and initiates an adhesion cascade that results in transendothelial migration (Ding et al., 2000). Inside the lymph node, pathogen-specific CTLs encounter antigen presenting cells (APCs) bearing pathogen-derived peptides and undergo clonal expansion as CD62L is replaced with adhesion molecules that facilitate migration to infected tissues.

Infected host cells produce large quantities of pro-inflammatory cytokines that orchestrate an immune response. Interleukin (IL)-2 and IL-12 promote formation of short-lived effector T cells (TEFF) that are programmed for an immediate lytic response (Chowdhury et al., 2011). Some TEFF cells express Killer Cell Lectin Like Receptor G1 (KLRG1), which is a membrane-bound adhesion molecule with an immunoreceptor tyrosine-based inhibitory motif (ITIM) in the cytoplasmic tail (Ito et al., 2006; Tessmer et al., 2007). Although most CTLs transiently express KLRG1 during antigen stimulation (Herndler-Brandstetter et al., 2018; Joshi et al., 2007), terminally-differentiated TEFF cells maintain KLRG1 expression and do not convert to a memory phenotype (Dominguez et al., 2015). Most TEFF cells have limited capacity for long-term survival and disappear as the infection resolves, leaving residual populations of memory CD8 T cells in the circulation and peripheral tissues. Central memory CD8 T cells (TCM) enter the bloodstream and transit between secondary lymphoid organs (SLO), whereas peripheral tissues are colonized with tissue-resident memory CD8 T cells (TRM) that express CD69 (Mueller et al., 2013). How these different memory subsets augment immunity varies during infections with different types of pathogens. For instance, early recruitment of circulating memory CD8 T cells into the liver is important for control of malaria infection (Lefebvre et al., 2021). Conversely, IAV replicates in the lungs and immunity is facilitated by pulmonary TRM cells that reside near the site of pathogen-exposure (Wu et al., 2014). During respiratory infection, vigorous proliferation by circulating memory CD8 T cells is less desirable, as the lungs may be overwhelmed with massive numbers of TEFF cells (Wu *et al*., 2014). TGFβ is a suppressive cytokine that can be activated by viral neuraminidase, and dampen the inflammatory response in the lungs during infection with seasonal strains of IAV (Carlson et al., 2010). Continuous exposure to TGFβ is required for TRM cells to express of the cadherin-binding protein αEβ7 integrin (CD103) for enhanced retention in the epithelial layer (Cepek et al., 1994; Teraki and Shiohara, 2002).

Cytokines that belong to the TGF family elicit cellular responses via both canonical and non-canonical signaling pathways. Canonical signals are mediated by a cascade of structurally-related molecules known as small (Sma) and mothers against decapentaplegic (Mad)-related (SMAD) proteins (Massague, 2012). SMAD4 is an adaptor that chaperones phosphorylated receptor-activated SMAD proteins (R-SMADs) into the nucleus to modulate gene expression. We previously used Cre-Lox recombination to determine how signaling via SMAD4 altered the surface markers on antiviral CTLs during IAV infection and found that very few SMAD4- deficient CTLs expressed KLRG1 or CD62L, while abnormal CD103 expression was detected on CTLs in the spleen (Hu et al., 2015a). These changes were unexpected, as CTLs displayed a reciprocal phenotype after the TGFβ receptor was ablated (Hu *et al*., 2015a). To further define how SMAD-dependent signaling pathways alter the migratory properties of pathogen-specific CTLs, we have used RNA-sequencing to identify genes that were either induced, or repressed, in the absence of SMAD4. Our data show that a novel SMAD4-dependent signaling pathway alters the expression levels of multiple transcription factors that are known to influence memory formation. Importantly, CD103 expression on TRM cells is induced by a conical SMAD-dependent signaling pathway during stimulation TGFβ, while an alternative SMAD4-dependent mechanism promotes formation of TEFF and TCM cells in the circulation.

## Materials and methods

### Mice and reagents

Mice were bred and housed at the University of Connecticut Health Center in accordance with institutional guidelines. Experiments were performed in accordance with protocols approved by the UCONN Health Institutional Animal Care and Use Committee (IACUC). Mice that express Cre-recombinase under the control of the distal-Lck promoter (dLck) were used to generate mice that lack R-SMAD2 (S2KO), R-SMAD3 (S3KO), co- SMAD4 (S4KO), EOMES (EKO) and TGFβ receptor II (TR2KO). S4KO and TR2KO were further crossed with OTI mice that express a transgenic antigen receptor specific for the SIINFEKL peptide presented on H-2K^b^.

Virus stocks were grown in fertilized chicken eggs (Charles River) and stored as described previously. Between 8 to 20 weeks (wks) after birth, anesthetized mice were infected intranasally (i.n.) with either 2x10^3^ plaque- forming units (PFU) X31-OVA, or 5x10^3^ colony-forming units (CFU) of recombinant *Listeria monocytogenes* expressing chicken ovalbumin (LM-OVA).

### Sample preparation for adoptive transfer and flow cytometry

Naïve CD8 T cells were isolated from SLO using Mojosort isolation kits (Biolegend, Dedham MA). Mice received 5x10^3^ congenically-marked donor cells by intravenous (i.v.) injection given 48 hours (hrs) before infection. To identify bloodborne CTLs by intravascular staining, mice were injected with 1μg anti-CD8β in 200ul PBS and sacrificed 5 minutes later. For flow cytometry, chopped lung tissue was incubated at 37°C for 90 minutes (mins) in RPMI with 5% fetal bovine serum (FBS) and 150 U/ml collagenase (Life Technologies, Rockville, MD, USA). Nonadherent cells were enriched on Percoll density gradients (44/67%) spun at 1200*g* for 20 min. Lymphocytes were incubated with antibodies that block Fc-receptors (15 mins at RT). Antigen-experienced CD8 T cells were identified using high CD11a/CD44 expression and divided into subsets with antibodies specific for CD103, KLRG1, CD127 and CD62L. For intracellular staining, lymphocytes were analyzed using True Nuclear transcription factor buffer (Biolegend, Dedham-MA). Permeabilized cells were stained with antibodies specific for T-bet and EOMES. To monitor cell-proliferation, cells were labeled with carboxyfluorescein succinimidyl ester (CFSE) dye before transfer to C57BL/6 mice. Alternatively, mice received a single injection of 5-Bromo-2-deoxyuridine (BrdU) by i.p. injection at 30dpi and donor cells were analyzed 3-6hrs later, using BrdU analysis Kits (BD Pharmingen). Stained cells were analyzed using a BD LSRII with FACSDiva V8.0 (BD Biosciences) and processed using Flowjo® and Graphpad Prism software. To visualize intracellular proteins, fixed cells were permeabilized using 90% ice cold methanol for intracellular staining. CTLs were imaged at 60X normal magnification using an Amnis ImagestreamX MKII flow cytometer. Similarity scores were calculated using Amnis Ideas V6.2 (Luminex) software.

### Cell culture

Naïve CD8 T cells were grown in RPMI with 10% FBS, L-glutamine, β-mercapthoethanol, sodium pyruvate, HEPES, and antibiotics. For T cell receptor (TcR) stimulation, CTLs were stimulated with plate-bound anti- CD3/CD28 (1μg/ml) and rIL-2 (20 U/ml). After 48hrs, CTLs were transferred to new wells for cytokine stimulation without TcR stimulation. Replicate wells were supplemented with rIL-2 (20 U/ml), ALK5 inhibitor (SB431542, 10μM/ml), and TGFβ (10 ng/ml).

### RNA-sequencing and gene set enrichment analysis

Messenger-RNA was extracted from purified CTLs using RNAeasy Plus Mini Kits (Qiagen, Hilden Germany) and sequenced by Otogenetics Corporation (Atlanta, GA). RNA-sequences from OTI-S4KO and OTI-Ctrl cells are available in the GEO database (accession # GSE151637). Data were analyzed using Tophat2, Cufflinks and Cuffdiff software (DNAnexus, Mountain View, CA). Transcriptional changes greater than two-fold (Log2 fold change >1 or <-1, p<0.05, false discovery rate<0.05) were used to select genes for Ingenuity pathway (Qiagen, Hilden Germany) and gene set enrichment analysis (GSEA) (Broad Institute, San Diego CA) (Subramanian et al., 2005). Differentially expressed genes were compared with published datasets, using GEO accession #s GSE39152 (Wakim et al., 2012), GSE107281(Milner et al., 2017), GSE70813 (Mackay et al., 2016), and GSE125471 (Nath et al., 2019). Normalized enrichment scores (NES) and false discovery rates (FDR) were visualized using ggplot2 package in R.

### RNA isolation for quantitative PCR analysis

Total RNA was extracted using Quick RNA MiniPrep Plus kit (Zymo Research). The quantity and quality of the RNA was analyzed using NanoDrop. Total RNA was reverse transcribed using M-MuLV reverse transcriptase (New England Biolabs) following the manufacturer’s instruction. Quantitative real-time PCR was performed using 2X SYBR green master mix (Bimake). Relative gene expression was determined using ddCt method with Ribosomal protein L9 (rpl9) as reference. Sequences for primers are shown in table 1.

### Retroviral transduction

A murine stem cell virus (MSCV)-IRES-GFP (pMig)-EOMES vector (RV-EOMES-GFP) was used for ectopic gene expression in activated CD8 T cells (Pearce et al., 2003). Viruses were packaged in human embryonic kidney (HEK) 293 cells and viral supernatants were processed using Retro-X concentrators (Takara Bio. USA Inc). Naïve CD8 T cells were purified by negative-selection and stimulated *in vitro* with plate-bound anti- CD3/CD28. After 48hrs, activated CD8 T cells were spin-infected (1800*g*, 60 min, 30°C) using concentrated supernatants and polybrene (1 μg/ml). After 3hrs, the washed cells were cultured in complete RPMI.

### Statistical analysis

To evaluate variability between biological replicates, P values were calculated using two-way ANOVA followed by Tukey’s multiple comparison tests, using Graphpad Prism®. Horizontal lines indicate comparisons between samples. For *in vitro* experiments, variability between technical replicates was evaluated using Student’s *t* Tests calculated using Graphpad Prism®. Experiments were repeated 2 or 3 times, based on outcomes.

## Results

### SMAD4 alters homing receptor expression in the absence of TGFβ

Several groups used CreLox recombination to prevent SMAD4 (S4KO), or TGFβ receptor II (TR2KO) expression in peripheral CD8 T cells (Zhang et al., 2005). For these studies, Cre is expressed under the control of the distal Lck (dLck) promoter to induce DNA at a late stage during thymic development (ref). When we analyzed homing-receptors during IAV infection, we found that antiviral CTLs displayed very different phenotypes in the absence of either SMAD4 or TGFβRII (Cao *et al*., 2015; Hu *et al*., 2015a). Since SMAD4 and TGFβRII are components of an interconnected signaling network, we intercrossed S4KO and TR2KO mice to produce CTLs that lack both molecules (S4TR2-DKO). For transfer experiments, the mutant mice were further crossed with OTI mice that expressed a transgenic antigen-receptor that is specific for a peptide (SIINFEKL) derived from chicken ovalbumin (OVA) presented on H-2K^b^ (Hogquist et al., 1994). The mice were infected with recombinant pathogens (X31-OVA and LM-OVA) that express a peptide derived from chicken ovalbumin (Jenkins et al., 2006; Pope et al., 2001).

To analyze homing-receptor expression during infection, congenically-marked OTI-Ctrl (lack Cre), OTI- TR2KO; OTI-S4KO and OTI-S4TR2DKO cells were transferred to B6 mice. After 48hrs the mice were infected with pathogens that elicit different inflammatory responses. IAV was used for a mild inflammatory response, as neuraminidase activates TGFβ (Schultz-Cherry and Hinshaw, 1996). Other mice were infected with *Listeria Monocytogenes* (LM) to generate TEFF cells that express KLRG1 (Plumlee et al., 2015). On different days post infection (dpi), donor cells in the lungs and spleens were analyzed for CD103, KLRG1, CD62L and CD127 expression. Heat maps show donor cells in the lungs and spleens after infection with X31-OVA **(Fig. 1A),** and in the spleens after infection LM-OVA **(Fig. 1B).** Statistical comparisons are shown in the supplementary data **(Supplementary Tables 1** and **2)**. The percentages of control CTLs that expressed KLRG1 and CD62L varied during the two infections, indicating exposure to different inflammatory conditions. The majority of OTI-TR2KO cells expressed KLRG1 at 8dpi and CD62L began to reappear by 30dpi. In contrast, smaller percentages of OTI-S4KO cells expressed KLRG1 and CD62L during both infections, indicate a defect during formation of TEFF and TCM cells. Importantly, OTI-S4KO and OTI-S4TR2-DKO cells displayed very similar phenotypes at all time points analyzed. These experiments show that SMAD4 alters homing-receptor expression via a TGFβ independent pathway. We used previously used CFSE and BrdU incorporation to measure cell-proliferation during IAV infection and found OTI-S4KO and OTI-S4Ctrl cells proliferated at similar rates (Hu *et al*., 2015b). When OTI-S4TR2DKO and OTI-S4TR2Ctrl cells were analyzed by the same approach, there was also no difference in cell-division **(Supplementary Fig. S1A).** These data indicate the SMAD4-ablation using dLck-Cre does not impede T cell priming.

**Figure 1.**
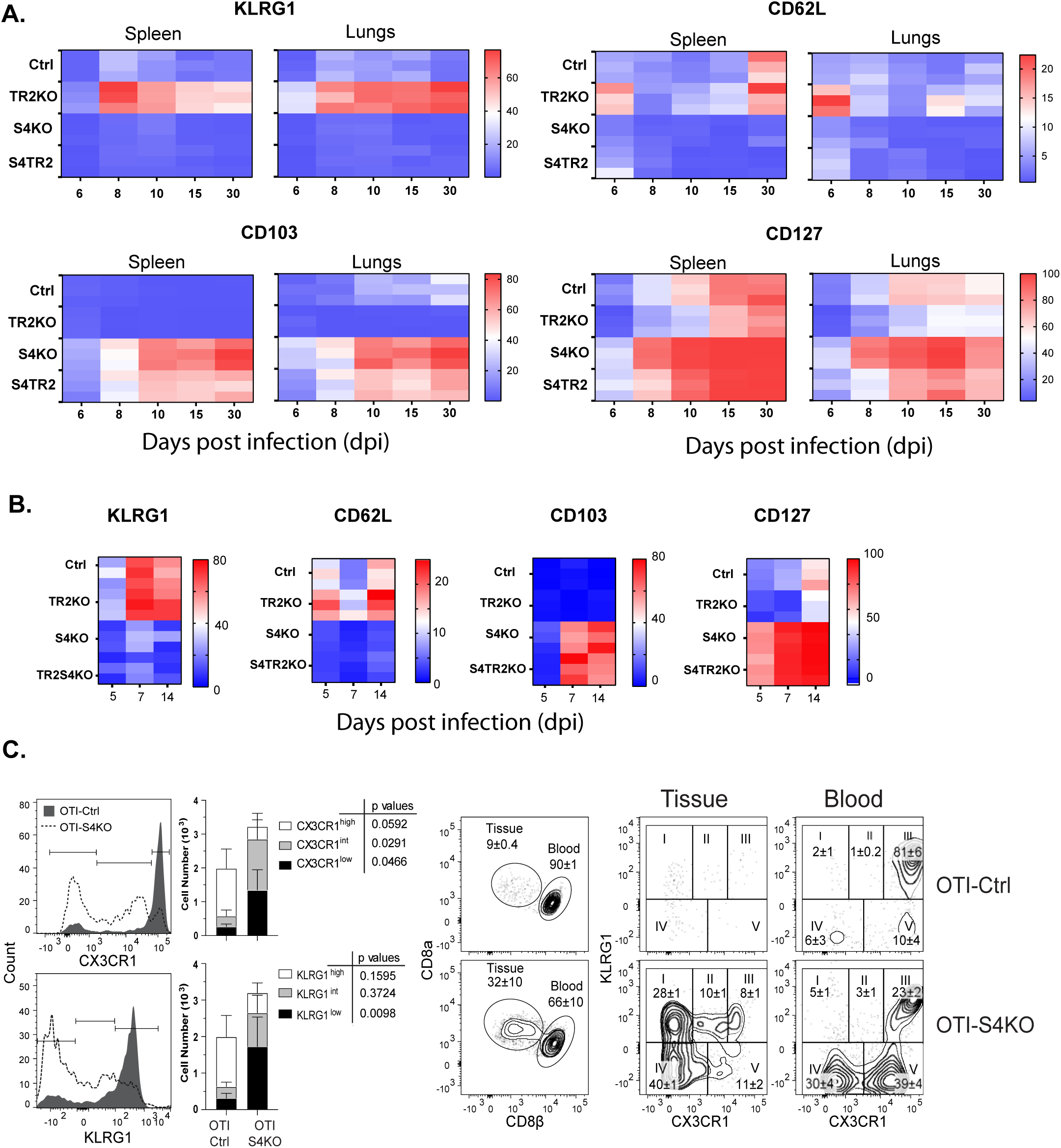
SMAD4-ablation alters homing receptor expression without TGFβ receptor II. **A&B).** Naïve donor cells from OTI-Ctrl, OTI-TR2KO, OTI-S4KO and OTI-S4TR2-DKO were transferred to B6 mice before infection. Heatmaps show percentages of donor cells that expressed KLRG1, CD103, CD62L and CD127. Data are means +SD (n=3/group). Two independent experiments gave similar results. **A).** CTLs in the spleens after infection with LM-OVA **B)** CTLs in the lungs and spleens after infection with X31-OVA **C).** B6 mice received mixed donor cells (OTI-S4KO and OTI-S4Ctrl) before infection with X31-OVA. Antibodies (CD8β) were injected 5 mins before sacrifice. Donor cells in lungs were analyzed for KLRG1 and GFP (CX3CR1) expression at 34dpi. Histograms (percentages) and bar graphs (cell numbers) show donor cells from OTI-S4KO (dashed) and OTI-S4Ctrl mice (continuous line). Contour plots show donor cells in the bloodstream and lung tissue.

Fractalkine (CX3CL1) is an unusual chemoattractant with adhesive properties (Ostuni et al., 2020), and alters CD8 T cell migration. The receptor (CX3CR1) is expressed at high levels on CTLs that localized in the vasculature(Nishimura et al., 2002). Reporter mice were used to analyze fractalkine receptor (CX3CR1^GFP^) expression during infection and showed that terminally-differentiated TEFF cells express both KLRG1 and CX3CR1 at high levels (Gerlach et al., 2016; Jung et al., 2000). Since a small percentage of OTI-S4KO cells expressed KLRG1 during infection with LM-OVA, we investigated whether CX3CR1 was regulated via SMAD4. To generate donor mice for transfer experiments, the CX3CR1^GFP+^ reporter was bred with OTI-S4KO mice. To analyze the CTL response, OTI-Ctrl and OTI-S4KO cells were mixed (1:1 ratio) and transferred to B6 mice before infection with LM-OVA. After 30 days, donor cells in the lungs and spleens were analyzed for KLRG1 and GFP expression **(Fig 1C and supplementary Fig. S1D)**. Fluorescently-conjugated antibodies (CD8β) were injected 5 mins before sacrifice to distinguish CTLs in the blood vessels and peripheral tissues. The majority of OTI-Ctrl cells (top row) expressed KLRG1 and GFP at high levels (histogram and bar graph) and were located inside the blood vessels (contour plots), whereas OTI-SKO cells expressed KLRG1 and GFP at intermediate or low levels, and included some cells that were located outside of the vasculature. Low numbers of CTLs that expressed KLRG1 and CX3CR1 at high levels indicated that SMAD4 was a catalyst for terminal-differentiation.

Rodents lose large percentages of their body weight during IAV infection and the rate of recovery to normal size can be used to assess the severity of disease. After IAV infection, S4KO mice experienced more severe disease than the controls (Hu *et al*., 2015a). Since OTI-S4KO and OTI-S4TR2-DKO cells display similar phenotypes during infection, we infected S4TR2-DKO and S4TR2-Ctrl mice with X31-OVA and recorded weight changes **(Supplementary Fig. S1B).** As expected, the S4TR2-DKO mice lost more weight and recovered from infection more slowly than the controls. After 30 days, the immune mice (and uninfected controls) were challenged with different strain of IAV (WSN-OVAI) **(Supplementary Fig. S1C)**. None of the immune mice experienced substantial weight loss after infection with WSN-OVAI, indicating that protection against reinfection was not dependent on SMAD4 or TGFβ.

### SMAD4 modifies homing-receptor expression independently of R-SMAD2/3

The TGF family includes two groups of cytokines, that signal via alternative branches of the SMAD cascade. TGFβ and activins signal via R-SMAD2/3, while bone morphogenic proteins (BMPs) and related growth differentiation factors (GDF) signal via R-SMAD1/5/8 (Fink et al., 2003; Takimoto et al., 2010). SMAD4 facilitates signaling by both groups of cytokines and chaperones complexes of phosphorylated R-SMADs into the nucleus for gene regulation (Massague et al., 2005). There is some redundancy within the SMAD signaling network, since experiments with cancer cells show that heterodimers comprised of R-SMAD2 and R-SMAD3 can enter the nucleus without SMAD4, after TGFβ stimulation (Fink *et al*., 2003; Li et al., 2008). In CTLs the TGFβ receptor phosphorylates R-SMAD3 to induce CD103 expression (Yang et al., 1999). Since our data indicate that SMAD4 regulates CTLs independently of TGFβ, we used cultured CTLs to investigate whether signaling via R-SMAD2/3 influenced the phenotype of S4KO cells during cytokine stimulation. dLck-Cre mice were crossed with mice that carry flox sites in R-SMAD2 (Ju et al., 2006) and R-SMAD3(Li *et al*., 2008), to produce S2KO (lack R-SMAD2); R- S3KO (lack R-SMAD3), and S23DKO (lack both molecules) mice. S3KO were also bred with S4KO mice to produce S34DKO (lack R-SMAD3 and SMAD4). For each line, the respective Cre-deficient littermates were used as controls.

To analyze phenotypic changes during cytokine stimulation, Naïve CD8 T cells were isolated from SLO by negative-selection and activated with plate-bound anti-CD3/CD28 plus recombinant IL-2 (20u/ml). After 48hrs the remaining CTLs were washed and cultured in fresh medium without TcR stimulation (48hrs).

Replicate wells were supplemented with either rIL-2 alone, or rIL-2 plus exogenous TGFβ (10ng/ml). Selected wells were supplemented with an ALK5 inhibitor (SB) to block responses to any TGFβ in the media. At 96hrs after TcR stimulation, the cultured CTLs were analyzed for CD103 **(Fig 2A)** and CD62L expression **(Fig 2B)**.

**Figure 2.**
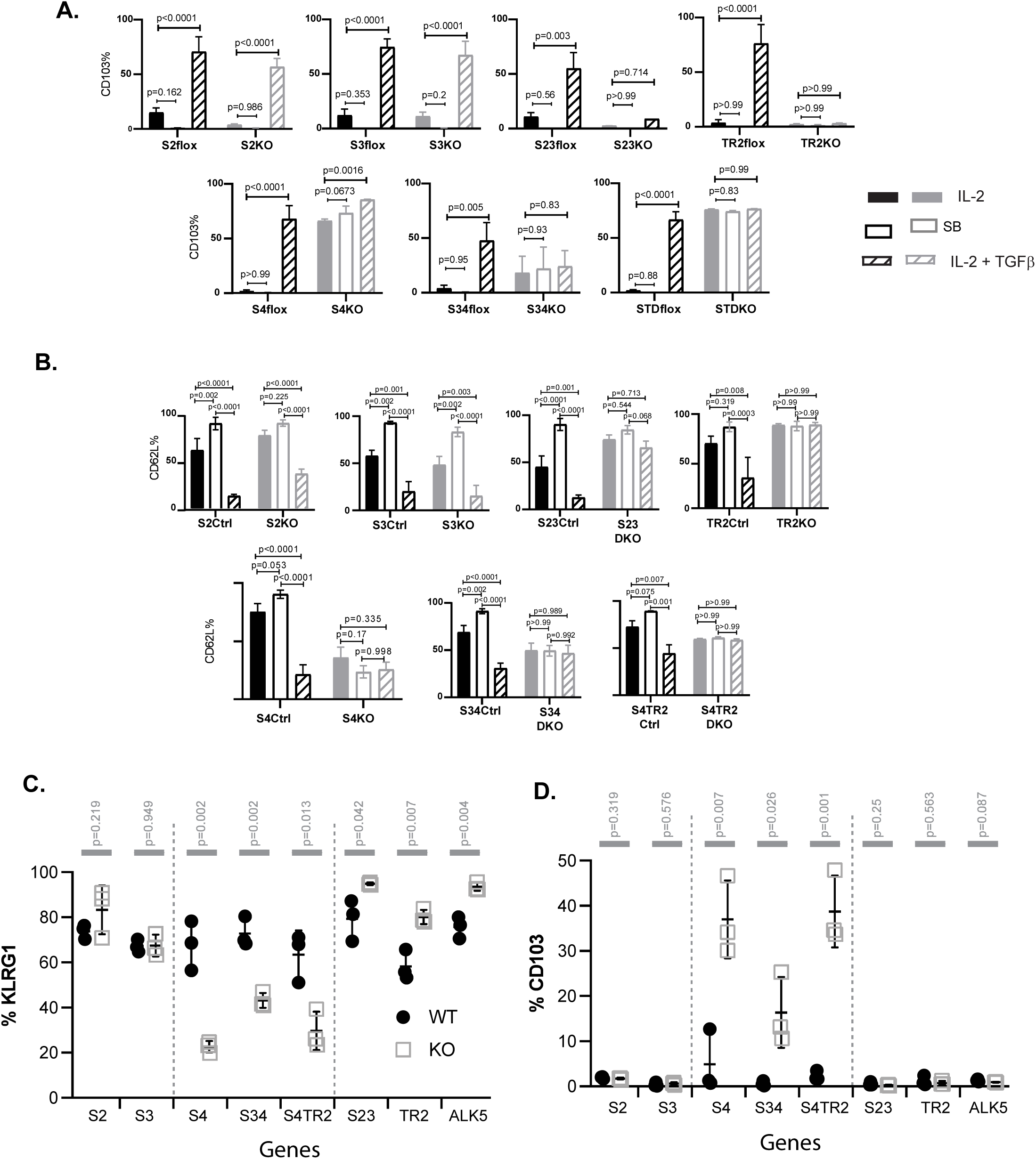
R-SMAD2/3 do not alter the phenotype of SMAD4-deficient CTLs. **A and B)** Naive CD8 T cells were activated *in vitro*. Selected wells were supplemented with SB431542 (10μM). After 48hrs, CTLs were transferred to new wells for secondary cytokine stimulation. Bars graphs are CTLs stimulated with rIL-2 alone (no fill); rIL-2 plus SB431542 (grey fill) and rIL-2 plus TGFβ (hatched). Data are means + SD (3 mice per group). P values were calculated using Student T tests. Two independent experiments produced similar results. **A)** Percentages of CTLs that expressed CD103. **B)** Percentages of CTLs that expressed CD62L **C and D)** Different strains of genetically-modified mice were infected with LM-OVA. OVA-specific CTLs were analyzed with MHCI tetramers at 8 dpi. Data are means + SD (3 mice per group). P values were calculated using Student T tests. Two independent experiments produced similar results. **C)** Tetramer+ CTLs analyzed for KLRG1 expression **D)** Tetramer+ CTLs analyzed for CD103 expression

In normal mice, naïve CD8 T cells down-regulate CD103 during antigen stimulation and expression returns when TRM cells settle in tissues that contain TGFβ(Mackay et al., 2015). Consistently, the control CTLs did not express CD103 when cultured with rIL-2 (plus and minus ALK5 inhibitor), but expression was induced by TGFβ **(Fig 2A).** This pattern of CD103 expression was not altered on S2KO and S3KO mice. Conversely, S23DKO and TR2KO cells lacked CD103 expression under all conditions analyzed. These data show that, in the absence of R-SMAD2, R-SMAD3 cooperates with SMAD4 to upregulate CD103. Sloan Kettering Institute (SKI) protein is a nuclear proto-oncogene that inhibits TGFβ signaling through interactions with SMAD proteins (Sun et al., 1999). A prior report found that SMAD4 interacts with SKI to create heterodimers that repress *Itgae* expression (αE integrin) in CTLs(Wu et al., 2021). Consistently, we found that newly activated S4KO cells expressed CD103 at higher levels than the control cells during stimulation with rIL-2. Importantly, we observed a significant increase in CD103 expression when S4KO cells were stimulated with TGFβ. The S34KO and S4TR2-DKO cells also expressed CD103 at intermediate levels after culture in rIL-2, but there was no response to TGFβ. These data show that TGFβ requires R-SMAD2/3 to upregulate CD103 in the absence of SMAD4. To confirm this observation, we used imaging flow cytometry to visualize R-SMAD2/3 expression in OTI-S4KO and OTI-S4Ctrl cells at 8dpi with X31-OVA **(Supplementary Fig. S2A)**. As expected, SMAD4- ablation did not influence the portion of R-SMAD2/3 that entered the nuclei.

The *in vivo* data show that most OTI-S4KO cells lacked CD62L expression at 30dpi, indicating a defect during formation of TCM cells. Since TGFβ inhibits TCM formation(Takai et al., 2013), we investigated whether R- SMAD2/3 controlled CD62L expression *in vitro* **(Fig 2B)**. To address this question, CTLs were activated and stimulated with cytokines as previously. Between 50-60% of the control CTLs expressed CD62L during culture with rIL-2 and the percentages increased (∼100%) when the ALK5 inhibitor was used. Only 25% of control CTLs expressed CD62L when the cultures contained TGFβ. After SMAD4 was ablated, 50% of S4KO, S34DKO and S4TR2DKO cells expressed CD62L during culture with IL-2 and there were no changes when the ALK5 inhibitor, or exogenous TGFβ was used. The TR2KO and S23KO cells expressed CD62L at high levels under all conditions analyzed. These data indicate that SMAD4 has dual functions and down-regulates CD62L in conjunction with TGFβ, and also induces CD62L expression independently of TGFβ.

Since recombinant cytokines do not induce KLRG1 expression on cultured CTLs (Robbins et al., 2003), we infected different strains of mutant mice with LM-OVA and analyzed TEFF in spleens at 8dpi. OVA-specific CTLs were analyzed with MHCI tetramers. The graphs show percentages of CTLs that expressed KLRG1 **(Fig 2C)** and CD103 **(Fig 2D)**. Between 50-90% of the control CTLs expressed KLRG1, and these numbers did not change in the absence of either R-SMAD2 (S2KO); R-SMAD3 (S3KO). KLRG1 was expressed on 80-90% of CTLs that lacked R-SMAD2/3 (S23DKO), or the TGFβ receptor (TR2KO and ALK5KO), whereas smaller percentages of CTLs expressed KLRG1 in the absence of SMAD4 (S4KO, S34DKO and S4TR2DKO). CD103 was only expressed on CTLs that lacked SMAD4 **(Fig 2D)**. These data show that SMAD4 promotes formation of terminally-differentiated TEFF cells via a mechanism that does not involve TGFβ or R-SMAD2/3.

### The transcriptional profile of SMAD4-deficient CTLs is similar to TRM cells

Although TGFβ is an important regulatory factor during CTL development, relatively little is known about the function of SMAD4 during transcriptional programming of activated CTLs. To address this question, we used RNA-sequencing to compare the transcriptional profiles of SMAD4-deficient and control CTLs during IAV infection. Early effector (EE) cells are partially-differentiated CTLs that have not committed to a classical SLEC/MPEC phenotype, and can be identified using high CD44/CD11a expression in the absence of KLRG1, CD127, and CD103 (Plumlee *et al*., 2015). To analyze phenotypically similar populations of antigen-specific CTLs, naïve OTI-S4KO and OTI-S4Ctrl cells were transferred to B6 mice before infection with X31-OVA and EE cells were isolated from the spleens 6dpi. Samples were analyzed in triplicate and mRNA-sequences were registered with the GEO repository (Accession # GSE151637). Gene set enrichment revealed many differentially-expressed genes **(Fig. 3A)**. Notably, OTI-S4KO cells expressed EOMES (*Eomes*) and CD62L (*Sell*) at lower levels than the controls, while *Itgae* was upregulated **(Fig. 3B)**. As large numbers of mucosal TRM cells express CD103, we used GSEA to compare the transcriptional profile of OTI-S4KO cells with a previously published data derived from TRM cells in the small intestine(Milner *et al*., 2017) **(Fig 3C)** and other tissues **(Fig 3D)**. Many of the core signature genes that are associated with TRM cells were upregulated in OTI- S4KO cells. We found similar correlations between OTI-S4KO cells and TRM cells isolated from the brain and liver, as well as CTLs that were stimulated with TGFβ *in vitro* (Mackay *et al*., 2016; Milner *et al*., 2017; Nath *et al*., 2019; Wakim *et al*., 2012). These correlations indicate that early signaling via SMAD4 impedes differentiation towards a resident memory phenotype.

**Figure 3.**
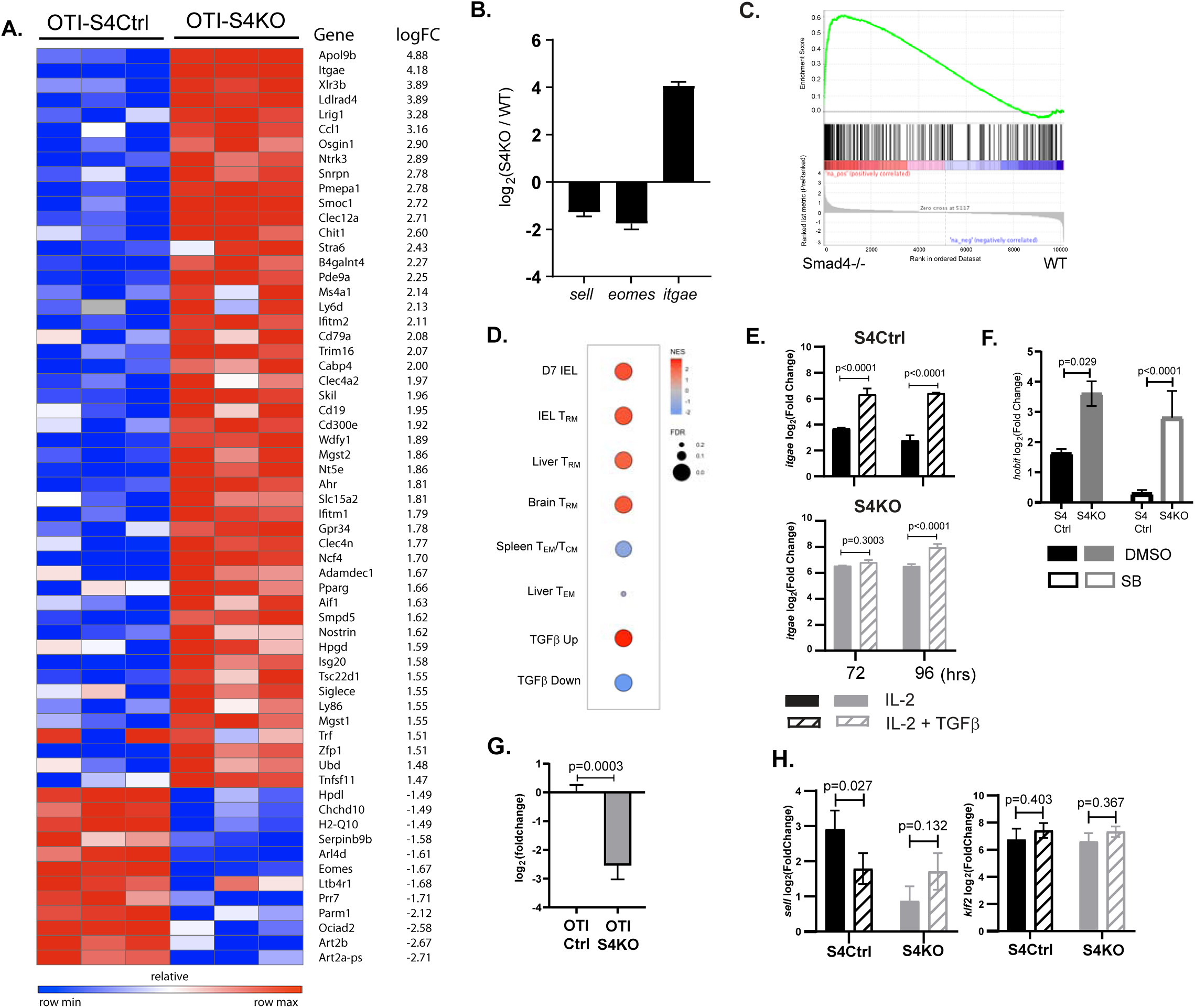
The transcriptional profile of SMAD4-deficient CTLs resembles a TRM phenotype. **A-C)** OTI-S4Ctrl and OTI-S4KO were transferred to B6 mice before infection with X31-OVA. Early effector (EE) cells were analyzed 6dpi, using RNA isolated from pools of three spleens. Samples were analyzed in triplicate (Total n=9/group). **A).** Differentially expressed genes were identified by enrichment analysis and ranked according to expression in OTI-S4KO cells. **B)**. *Eomes* and *Sell* genes were expressed at reduced levels in OTI-S4KO cells, while *Itgae* was induced. **C&D).** The transcriptional profile of OTI-S4KO cells was compared with published data sets. Comparisons are IEL 7dpi (GSE107395), IEL 30dpi, Liver T_RM_, T_EM_ and TCM (GSE70813), Brain TRM (GSE39152) and CD8 T cells activated *in vitro* in the presence and absence of TGFβ (GSE125471). **C)** Comparison with IEL analyzed 7dpi (GSE107395) **D)** Comparison with TRM cells in Brain Liver and in vitro CTLs. **E)** Cultured CTLs were analyzed for *Itgae* transcripts using qPCR analysis. Bars are cells stimulated with rIL-2 alone (Filled bars), and rIL-2 plus TGFβ (hatched bars). Samples were analyzed in triplicate. Two independent experiments gave similar results. **F)** Cultured CTLs were analyzed for *Hobit* transcripts using qPCR analysis. Bars are cells stimulated with rIL-2 alone (Filled bars), and rIL-2 plus ALK5 inhibitor (white bars). **G)** qPCR was used to measure *Klrg1* transcripts in OTI-S4KO and OTI-S4Ctrl cells at 8dpi with LMOVA. Samples were analyzed in triplicate. Two independent experiments gave similar results. **H)** Cultured CTLs were analyzed for *Sell* transcripts using qPCR analysis. Bars are CTLs stimulated with rIL-2 alone (Filled bars), and rIL-2 plus TGFβ (hatched bars). Samples were analyzed in triplicate. Two independent experiments gave similar results.

We next used qPCR to further confirm that SMAD4 alters *Itgae* expression *in vitro*. Naïve CD8 T cells were activated with plate-bound anti-CD3/CD28 plus recombinant IL-2 (20u/ml). After 48hrs, RNA was extracted from S4Ctrl cells to establish a baseline for gene expression. The remaining CTLs were cultured in fresh medium without TcR stimulation. Replicate wells were supplemented with either rIL-2 alone, or rIL-2 plus exogenous TGFβ (10ng/ml). RNA was extracted at 24hrs and 48hrs after cytokine stimulation (i.e. 72hrs and 96hrs after TcR stimulation, respectively) for qPCR analysis **(Fig. 3E)**. The graphs show relative changes in gene expression (log2 fold change) compared to the controls.

As expected, the S4KO cells produced more *Itgae* transcripts than S4Ctrl cells during culture with IL-2 and expression increased when TGFβ was added. Many transcription factors have important functions during CD8 T cell differentiation, including homolog of Blimp1 (Hobit) which promotes formation of TRM cells (Mackay *et al*., 2016; Parga-Vidal et al., 2021) and EOMES which is expressed in TCM cells (Intlekofer et al., 2005) and down-regulated by TGFβ as TRM cells settle in peripheral tissues(Mackay *et al*., 2015). Since our sequencing data indicated that SMAD4-ablation promoted a TRM phenotype, we used qPCR to measure *Hobit* transcripts. Naïve S4KO and S4Ctrl cells were activated with antibodies (48hrs) and cultured with rIL-2 (48hrs) in the presence and absence of the ALK5 inhibitor. Samples were analyzed by qPCR at 96hrs after TcR stimulation **(Fig 3F)**. We found that S4KO cells produced more *Hobit* transcripts than S4Ctrl cells during culture with IL-2. The numbers of *Hobit* transcripts in the S4Ctrl cells decreased when the ALK5 inhibitor was used, while expression in S4KO cells did not change. These data indicate that *Hobit* is induced by TGFβ and downregulated via SMAD4.

Because neuraminidase can activate TGFβ, small percentages of OTI-Ctrl cells expressed KLRG1 during infection with X31-OVA **(Fig 1B).** Consistently, RNA-sequencing did not reveal significant differences in the numbers of *Klrg1* transcripts in OTI-S4KO and OTI-S4Ctrl cells at 6dpi. To generate large populations of TEFF cells, we transferred OTI-S4KO and OTI-S4Ctrl cells to B6 mice before infection with LM-OVA and analyzed donor cells in the spleen at 8dpi using qPCR **(Fig 3G)**. The data showed that OTI-S4KO cells produced fewer *Klrg1* transcripts than OTI-S4Ctrl cells during the TEFF response.

L-selectin (CD62L) is an adhesion molecule that is required for naïve and TCM cells to enter resting lymph nodes via HEV. Krupple-like factor 2 play a central role in transcriptional regulation of CD62L during the CTL response(Takada et al., 2011). Since our data indicate that CD62L is regulated via SMAD4(Cao *et al*., 2015; Hu *et al*., 2015b), we used cultured CTLs to measure *Sell* (CD62L) and *KLF2* transcripts after cytokine stimulation **(Fig. 3H).** Naive S4KO and S4Ctrl cells were activated (48hrs) and stimulated with rIL-2, or rIL-2 plus TGFβ (48hrs). qPCR analysis showed that S4KO cells produced fewer *Sell* transcripts than S4Ctrl cells during culture with rIL-2. S4Ctrl cells produced fewer *Sell* transcripts during stimulation TGFβ, while expression in S4KO cells did not change. These data indicate that CD62L is induced via SMAD4 to promote formation of TCM cells, and down-regulated by TGFβ to promote TRM development. The numbers of *Klf2* transcripts were not modified by SMAD4 or TGFβ.

### EOMES is cooperatively regulated by TGFβ and SMAD4

EOMES and T-bet are Tbox transcription factors with important functions during lineage-specification of pathogen-specific CTLs. Our sequencing data indicated that OTI-S4KO cells produced fewer *Eomes* transcripts than OTI-S4Ctrl cells during IAV infection **(Fig 3B)**. To confirm whether EOMES was regulate via SMAD4, we activated S4KO and S4Ctrl cells *in vitro* (48hrs) and measured *Eomes* transcripts after cytokine stimulation using qPCR **(Fig 4A).** The numbers of *Eomes* transcripts in S4Ctrl cells transiently decreased during culture with rIL-2 (72hrs after TcR stimulation) and returned to baseline at 96hrs. *Eomes* was downregulated by TGFβ at both time points. Importantly, the S4KO cells produced small numbers of *Eomes* transcripts during culture with IL-2 and did not respond to TGFβ. Since these data indicate that EOMES is positively-regulated via SMAD4 to support TEFF formation.

**Figure 4.**
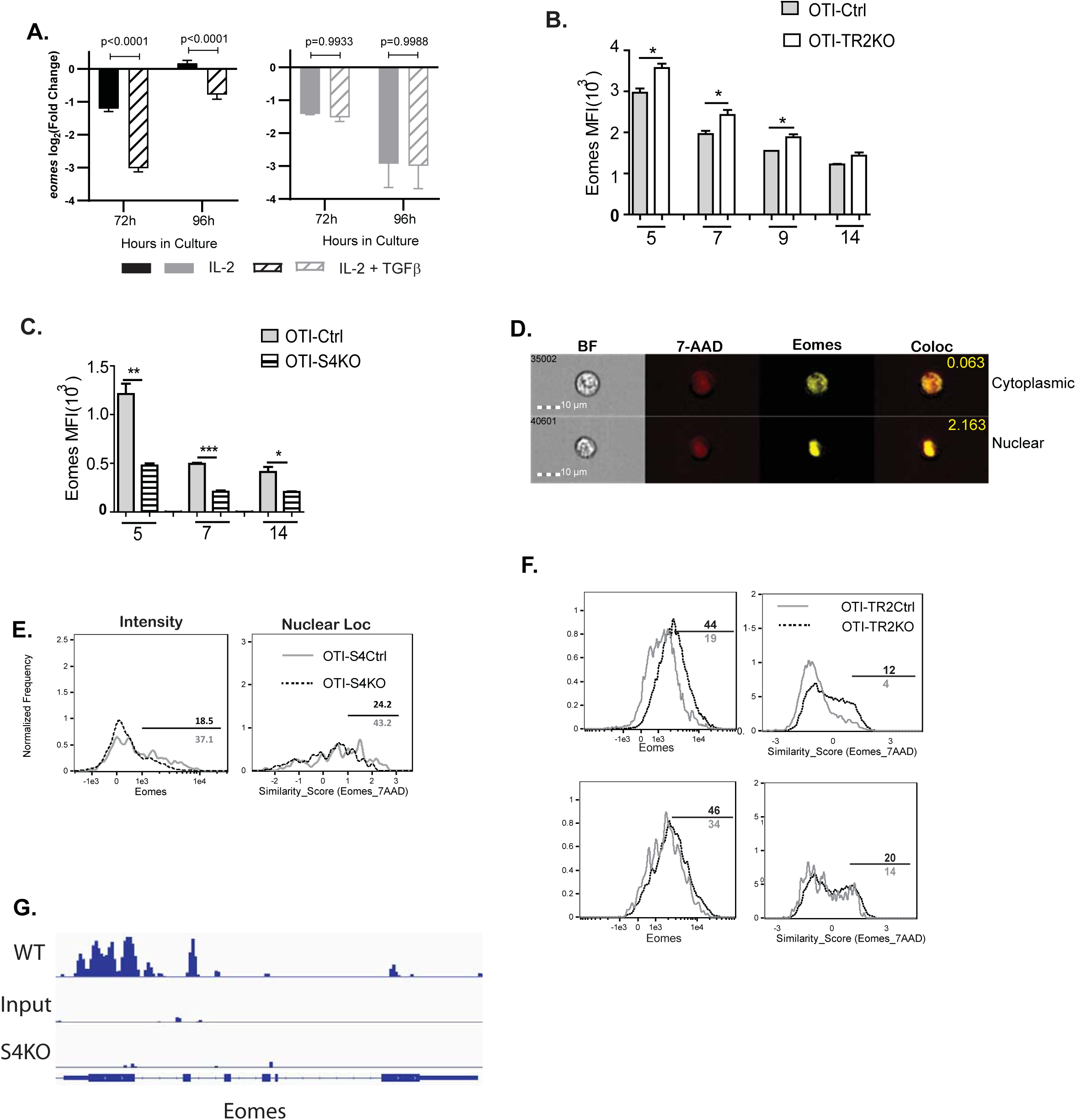
EOMES expression is induced via SMAD4 and downregulated by TGFβ. **A)** S4Ctrl (left) and S4KO (right) cells were activated *in vitro* and stimulated with cytokines. *Eomes* transcripts were measured at 96hrs after TcR stimulation by qPCR analysis. Bars show CTLs stimulated with rIL-2 alone (Filled bars), and rIL-2 plus TGFβ (hatched bars). Samples were analyzed in triplicate. Two independent experiments gave similar results. **B and C).** OTI-S4KO, OTI-S4Ctrl, OTI-S4TR2TKO and OTI-S4TR2Ctrl cells were transferred to B6 mice before infection with LM-OVA. On different days after infection, donor cells in the spleens were analyzed for EOMES expression by intracellular staining. Data are mean + SD (n=4/group). Two independent experiments gave similar results. **B)** EOMES expression (MFI) in OTI-S4KO (hatched) and OTI-S4Ctrl (grey fill) **C)** EOMES expression (MFI) in OTI-TR2KO (white bars) and OTI-TR2Ctrl (grey fill). **D-F).** OTI-S4Ctrl and OTI-S4KO were transferred to C57BL/6 mice before infection with LM-OVA. Imaging flow cytometry was used to analyze donor cells in the lungs at 8 dpi. Data are means + SD, n=3. **D).** Representative staining for EOMES (yellow) and 7-AAD (red) to identify the nuclei. **E)**. Histograms show EOMES intensity and similarity scores for OTI-S4Ctrl (grey line) and OTI-S4KO (dashed line) cells. Similarity scores were determined using IDEAS software. Higher similarity scores indicate that EOMES has translocated into the nucleus. **F)** Histograms show EOMES intensity and similarity scores for OTI-TR2Ctrl (grey line) and OTITR2KO (dashed line) cells. **G).** Published Chip-Seq data from *in vitro* activated CTLs were analyzed for SMAD4 binding-sites. The graph shows multiple SMAD4-binding sites in the *Eomes* locus.

We next used transfer experiments to monitor changes in EOMES and Tbet expression during infection. Naïve OTI-S4KO; OTI-S4Ctrl; OTI-TR2KO and OTI-TR2Ctrl mice were transferred to B6 mice 48hrs before infection with LM-OVA. On different days post infection (dpi), donor cells in the spleens were analyzed for EOMES expression by intracellular staining. We found that TGFβ and SMAD4 altered EOMES expression in opposite directions. OTI-TR2KO cells expressed EOMES at higher levels than OTI-TR2Ctrl cells at all time point analyzed **(Fig. 4B)**, whereas OTI-S4KO cells expressed EOMES at lower levels than OTI-S4Ctrl cells **(Fig. 4C).** After infection with X31-OVA, the differences in EOMES expression were less pronounced, which may reflect the influence of virally-induced TGFβ **(Supplemental data Fig. 4A)**.

Others analyzed CTLs during LCMV infection and found that EOMES was concentrated in the nuclei of exhausted CTLs (McLane et al., 2021). We used imaging flow cytometry to determine whether EOMES entered the nuclei of OTI-S4KO cells during TEFF formation **(Fig. 4D-F).** Donor cells were transferred to B6 mice before infection with LM-OVA and the spleens were analyzed at 8dpi. The OTI-S4KO cells expressed EOMES at lower levels than OTI-S4Ctrl, and a smaller percentage of the protein was located in the nuclei where it could influence gene expression **(Fig 4E)**. We next compared EOMES expression in TR2KO and TR2Ctrl cells using a KLRG1 gate **(Fig 4F)**. When we analyzed KLRG1^+^ CTLs, we found no difference in EOMES expression for TR2KO and TR2Ctrl CTLs. A similar comparison of KLRG1^-negative^ cells showed that TR2KO cells expressed EOMES at higher levels than the controls **(Fig 4F)**. Since these data show that EOMES is regulated via SMAD4, we used published ChIP-seq data to determine whether the regulatory sequences for the *Eomes* gene include a SMAD4-binding site **(Fig 4G)**. ChIP-Seq data from *in vitro* activated CD8 T cells (Accession number GSE135533) was analyzed using integrative genomics viewer (IGV) software. The data show that the promoter region for the *Eomes* gene includes multiple SMAD4 binding sites.

### Ectopic EOMES expression alters the ratios of CTLs that express CD103 and CD62L

Prior studies indicate that EOMES must be down-regulated before TRM cells express CD103 (Mackay *et al*., 2015). Since our data show that OTI-S4KO express EOMES at reduced levels compared to controls, we reasoned that SMAD4 might indirectly influence homing-receptor expression by altering EOMES expression. To explore this possibility, we intercrossed S4KO and EOMES-deficient (EKO) mice to produce CTLs with both mutations (ES4KO). Cre-deficient littermates (ECtrl, S4Ctrl and ES4Ctrl) were used as controls. Naïve CD8 T cells were activated *in vitro* (48hrs) and stimulated with cytokines (48hrs) as previously. Selected wells were supplemented with the ALK5 inhibitor to prevent signaling via the TGFβ receptor. CTLs were analyzed for CD103 **(Fig. 5A)** and CD62L expression **(Fig. 5B)** at 96hrs after TcR stimulation.

**Figure 5.**
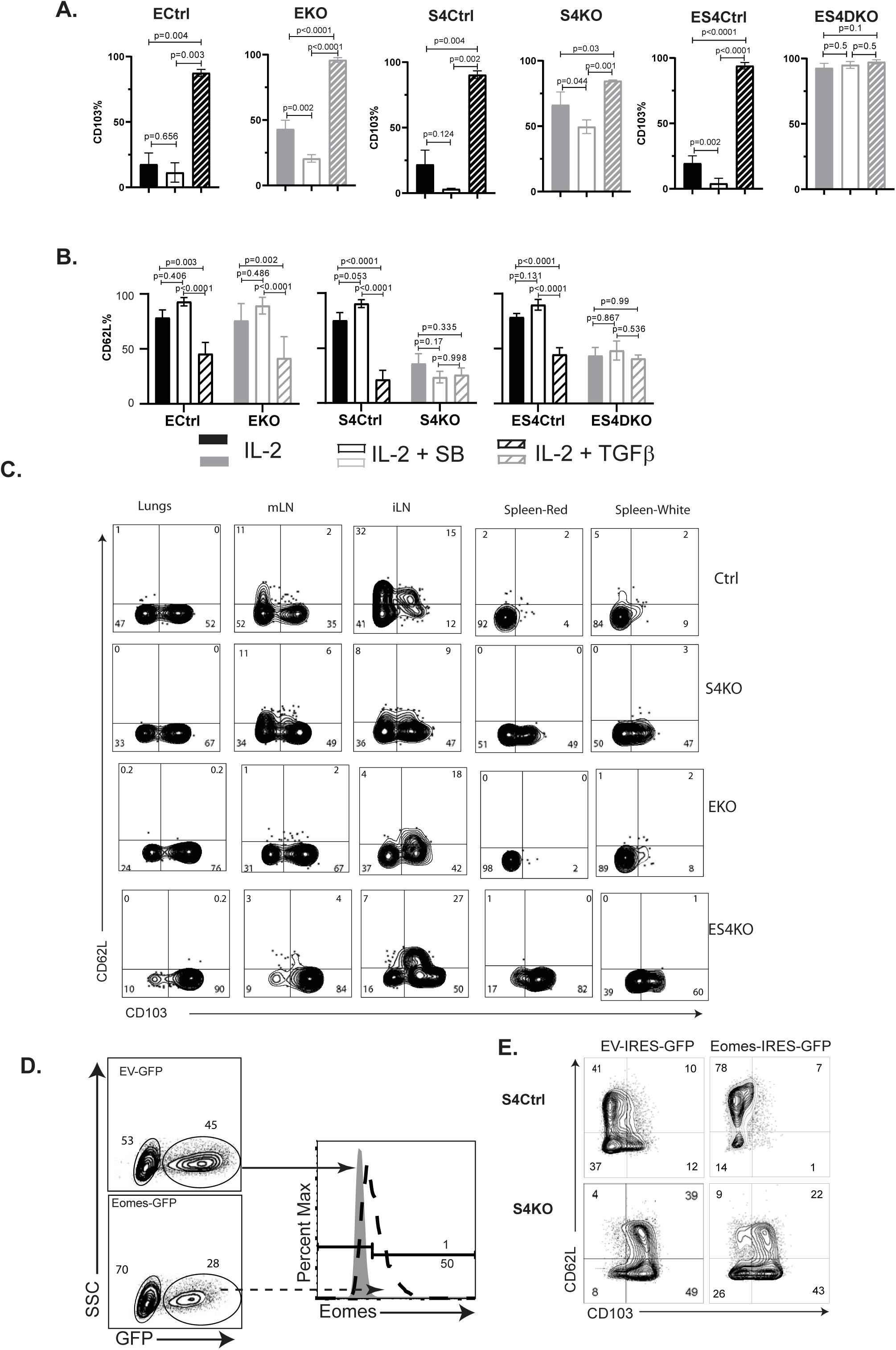
Ectopic EOMES expression facilitates TCM formation. A and B) Naïve CD8 T cells were activated *in vitro* (48hrs) and stimulated with cytokines (48hrs) *in vitro*. Bars graphs show CTLs stimulated with rIL-2 alone (no fill); rIL-2 plus SB431542 (grey fill) and rIL-2 plus TGFβ (hatched). Data are means + SD (3 mice per group). P values were calculated using student’s T tests. Two independent experiments produced similar results. A) Percentages of CTLs that expressed CD103 B) Percentages of CTLs that expressed CD62L C) S4KO, EKO, ES4KO mice were infected with X31-OVA. Anti-viral CTLs were analyzed with MHCI tetramers at 30dpi. Anti-CD8 antibodies were injected 5 mins before sacrifice. Contour plots show OVA-specific CTLs analyzed for CD103 and CD62L expression. D-E). EKO, S4KO and S4Ctrl cells were activated *in vitro* (48hrs) and transduced with a retroviral vector encoding EOMES (Eomes-IRES-GFP), or the empty vector control (EV-IRES-GFP). GFP+ cells were analyzed by flow cytometry at 96hrs after TcR stimulation. D). Overlaid histograms show EKO cells transduced with Eomes-IRES-GFP (grey shading), or EVIRES-GFP (dashed line). E) Contour plots show GFP+ S4KO and S4Ctrl cells analyzed CD62L and CD103 expression. Data are means + SD (3 mice per group). Two independent experiments produced similar results.

As expected, the control cells did not express CD103 during culture with IL-2 and this marker was induced by TGFβ. The S4KO and EKO cells both expressed CD103 at intermediate levels during culture with rIL-2 and expression increased when TGFβ was added. Conversely, ES4KO cells expressed CD103 at high levels under all conditions analyzed. These data show that the *Itgae* gene was repressed by two different mechanisms. In wildtype mice, CD62L is expressed on naïve CD8 T cells, down-regulated during TEFF formation and re-expressed on TCM cells. We analyzed naïve CD8 T cells before culture and found similar levels of CD62L expression in each group of mice **(Supplemental Fig 5A).** The control CTLs downregulated CD62L at 48hrs after TcR stimulation *in vitro* **(Supplemental Fig 5B),** while a majority CTLs re-expressed CD62L during culture with rIL-2. The percentages of control CTLs that expressed CD62L increased when the ALK5 inhibitor was present and decreased when TGFβ was used. In contrast, only 50% of S4KO and ES4KO cells expressed CD62L under all conditions analyzed **(Fig 5B)**. These data show that antigen-experienced CTLs down-regulate CD62L in response to TGFβ, and that SMAD4 is required for CTLs to re-express CD62L during memory formation.

We also infected S4KO, EKO, and ES4KO mice (plus littermate controls) with X31-OVA, to analyze homing-receptor expression *in vivo* **(Fig. 5C)**. OVA-specific CTLs were analyzed at 30dpi using MHCI tetramers. Anti-CD8 antibodies were given by i.v. injection 3 mins before sacrifice, to stain CTLs in the bloodstream. As expected, the lungs and draining lymph nodes (MLN) from each group of mice contained CD103^+^ TRM cells. Abnormal CD103 was expression was detected on OVA-specific CTLs in the spleens from S4KO and ES4KO mice. Approximately 50% of OVA-specific CTLs in the ILNs from control mice re-expressed CD62L, indicating the presence of TCM cells. Smaller percentages of OVA-specific CTLs expressed CD62L in the ILNs of S4KO, EKO and ES4KO mice (17-22%), supporting the idea that signaling via SMAD4 promotes formation of TCM cells.

To explore whether ectopic EOMES altered the phenotype of S4KO cells, we used a murine stem cell virus (MSCV) vector that encodes green fluorescent protein (MSCV)-IRES-GFP (pMIG) to over-express EOMES in antigen-experienced CTLs *in vitro* (**Fig.5D**). The pMIG vector has been described previously (Pearce *et al*., 2003). Naïve S4KO and EKO cells were stimulated with anti-CD3/28 and rIL-2 (48hrs), then transduced with either EOMES-IRES-GFP, or empty vector control (EV-IRES-GFP). At 96hrs after TcR stimulation, the GFP^+^ CTLs were analyzed for CD103 expression by flow cytometry. EKO cells were used to verify that the RV-EOMES-GFP vector was functional (overlaid histograms) (**Fig.5D**). When EOMES was expressed ectopically, smaller percentages of S4KO and S4Ctrl cells expressed CD103, while the percentages of CD62L^+^ CTLs increased **(Fig. 5E and supplemental Fig S5C)**. These data show ectopic EOMES expression partially restored the phenotype of S4KO cells, and induced a shift toward a central memory phenotype.

## Discussion

Defining how antigen-specific CTLs differentiate into functionally-specialized subsets of pathogen-specific memory CD8 T cells is an important priority for vaccine development. Multiple models have been used to define the relationships between these phenotypically distinct cell populations (PMID: 34923638). Some studies suggest that antigen-specific CTLs follow a linear differentiation pathway by transitioning between subsets, while others favor a branching model of differentiation whereby multiple subsets of CTLs arise from separate pathways that diverge soon after antigen stimulation(Bannard et al., 2009). The stage at which activated CTLs commit to a specific phenotype remains controversial. The transcriptional programs that determine whether pathogen-specific CTLs localize in peripheral or lymphoid tissues are largely undefined.

Different approaches have been used to catalog the heritable traits of pathogen-specific CTLs. Some studies suggest that naïve CD8 T cells are not a uniform cell population. Limiting-dilution assays and fate-mapping experiments were used showed that naïve CD8 T cells give rise to mixed populations of TEFF and memory CD8 T cells(Plumlee *et al*., 2015; Stemberger et al., 2007). Another study found unequal inheritance of genetic markers during TEFF formation, indicating that CTLs are products of asymmetric cell-division(Chang et al., 2011; Pollizzi et al., 2016). Phenotypic markers also indicate that the differentiation program of memory CD8 T cells begins soon after antigen-stimulation, as the IL-7 receptor (CD127) is expressed CTLs that are predisposed toward memory formation and absent from TEFF cells that express KLRG1(Kaech et al., 2003).

Transcriptome analysis revealed the presence of TRM precursors in a pool of circulating TEFF cells(Kok et al., 2020). Although many studies support a model of diverging differentiation, the question of whether specialized traits are genetically-programed, or induced by external stimuli has not been resolved. To address this question, transfer experiments were used to track clonal populations of antigen-specific CD8 T cells during infections with different type of pathogens. The study showed that the population dynamics reflected the inflammatory environment that developed during infection(Plumlee et al., 2013). Large numbers of TEFF cells expressed KLRG1 during infection with *Listeria monocytogenes*, while this marker was less prevalent during infection with *Vesicular Stomatitis Virus*.

Several cytokines are known to influence the ‘fate decisions’ of newly activated CTLs, including interleukin-12 (IL-12) which supports TEFF formation(Mescher et al., 2006), whereas TGFβ has anti- inflammatory properties and encourages TRM cells to settle in peripheral tissues(Skon et al., 2013). *In vivo*, naïve CD8 T cells can be preconditioned to become TRM cells through interactions with APCs that activate TGFβ(Mani et al., 2019). We previously showed that signaling via SMAD4 was required for formation of TEFF and TCM cells, whereas TGFβ creates a bias toward TRM formation. Our current study extends this work by showing that SMAD4 and TGFβ are essential components of two distinct signaling pathways and that guide the fate decisions of newly activated CTLs to achieve alternative outcomes. These pathways control a key bifurcation in the differentiation pathway of antiviral CTLs, by regulating the same genes in opposite directions. We found that SMAD4-induced several genes that are expressed by CTLs in the circulation, including EOMES, KLRG1 and CD62L. These molecules are downregulated as TRM cells settle in peripheral tissues. The conclusion of our study is that SMAD4 acts independently of TGFβ to support commitment to TEFF and TCM lineages. A further observation of this work is that SMAD4 has multiple functions during regulation the *Itgae* gene (CD103), including gene suppression by SMAD4/SKI heterodimers(Wu *et al*., 2021), as well as via SMAD4-dependent induction of EOMES.

Memory CD8 T cells can be divided into subsets of based on their function, lifespan, and tissue localization. Cells in peripheral and lymphoid tissues express different combinations of homing-receptors, that are induced by integrated signals from the antigen receptor and costimulatory molecules, as well as cytokines that are released during infection. We previously showed that antiviral CTLs express several homing receptors that are regulated via SMAD4, including KLRG1, CD103 and CD62L. To further define how changes in homing receptor expression are coordinated as CTLs respond to infection, we used RNA-seq to compare gene expression in OTI-S4KO and OTI-Ctrl cells during IAV infection. The experiment revealed a pattern of differentially-expressed genes that resembled the transcriptional profile of TRM cells in other non-lymphoid tissues. Genes that are normally associated with formation of TEFF cells and TCM cells were expressed at reduced levels in OTI-S4KO cells, including Eomes, CD62L (*Sell*), and KLRG1 (*Klrg1*). Conversely, several genes that are normally associated with TRM cells were expressed at increased levels on OTI-S4KO cells including αE integrin (*Itgae*) and Hobit. Our data indicate that SMAD4 and TGFβ perform reciprocal functions during memory formation, as SMAD4 favors formation of circulating memory CD8 T cells and inhibits TRM development, whereas TGFβ inhibits TEFF formation and induces TRM cells.

TGFβ is a pleiotropic cytokine with anti-inflammatory properties and uses a network of interconnected signaling pathways to modulate gene expression. The canonical signaling pathway for TGFβ involves formation of R- SMAD2/3 heterodimers that enter the nucleus with together SMAD4 (ref). Experiments with genetically-modified cells reveal extensive redundancy within the SMAD signaling cascade. In cancer cells, R-SMAD2/3 heterodimers entered the nucleus without SMAD4 and changed gene expression (David et al., 2016; Fink *et al*., 2003; Levy and Hill, 2005). When SMAD4 is present, homodimers that are comprised of either R-SMAD2 or R-SMAD3 are sufficient for signaling transduction (Yagi et al., 1999). Since R-SMAD2 lacks a DNA-binding domain(Yagi *et al*., 1999), R-SMAD2 homodimers did not alter gene expression in macrophages or helper T cells during stimulation with TGFβ (Chung et al., 2012; Malhotra et al., 2010; Takimoto *et al*., 2010; Wang et al., 2013). Similarly, R-SMAD2 was not sufficient for TGFβ to induce CD103 expression on CTLs in the absence of either R-SMAD3 or SMAD4.

SMAD proteins modulate gene expression through interactions with numerous transcription factors. In CD8 T cells, SMAD4 forms complexes with SKI to directly repress the *Itgae* gene (Wu *et al*., 2021). Consistently, we found that exogenous TGFβ was not required for S4KO cells to express CD103 at intermediate levels after antigen stimulation. Others used retroviral vectors to ectopically express T-box transcription factors in activated CTLs during HSV infection and found reduced numbers of TRM cells in the skin when EOMES and T-bet were expressed at high levels(Mackay *et al*., 2015). Further evidence that these transcription factors interfere with formation of TRM cells includes the observation that reduced EOMES and T-bet expression facilitate TRM development (Laidlaw et al., 2014). To further understand how these transcription factors support memory formation, we investigated whether EOMES was regulated via SMAD4. Our experiments confirmed that EOMES was downregulated by TGFβ acting in combination with R-SMAD2/3. Conversely, we found that SMAD4 was required to maintain EOMES expression in activated CTLs. These experiments indicate that SMAD4 has multiple functions during regulation of CD103, as SMAD4/SKI heterodimers interact with regulatory sequences in the *Itgae* gene (Wu *et al*., 2021), while SMAD4 indirectly represses CD103 through EOMES induction. The influence of both mechanisms during CTL activation was evidenced by cumulative changes in CD103 expression in the absence of SMAD4 and EOMES.

T-box transcription factors have important functions during differentiation of both CD4^+^ and CD8^+^ T cells. For this study, we prevented gene expression in peripheral T cells using Cre expressed under the control of the distal Lck promoter. Although SMAD4-ablation caused extensive phenotypic changes in antiviral CTLs, the surface markers on CD4 T cells were largely unaltered (data not shown). Since CD8 T cells express EOMES at higher levels than CD4 T cells (Zhu et al., 2010), our data are consistent with the view that different mechanisms drive effector and memory formation in the CD8+ and CD4+ lineages.

Many transcription factors have important functions during CD8 T cell differentiation. The major players include Zeb1 and Zeb2, which are homologous proteins that serve opposing functions during memory formation and are inversely regulated by TGFβ. *Zeb1* expression increases in maturing memory CD8^+^ T cells during exposure to TGF-β, whereas *Zeb2* is down-regulated (Dominguez *et al*., 2015; Guan et al., 2018). Other studies indicate that DNA-binding inhibitors Id2 and Id3 also have reciprocal functions during fate determination of activated CTLs (Ji et al., 2011; Omilusik et al., 2018; Yang et al., 2011). Whether SMAD4 alters the expression levels of these transcription factors, or other epigenetic-regulators, during CD8 T cell differentiation needs to be further investigated.

## Acknowledgments

Cells were purified with the assistance of the Flow cytometry Core Facility at UCONN Health. This work was funded by NIH grant AI123864 (LSC and SK) and a postdoctoral career development award from the American Association of Immunologists (KC).

## Author Contributions

Conceptualization and investigation: K.C., J.S-R., Y.H., J.S.L., and Z.U. Formal analysis: B.M. Writing – Original Draft: K.C and L.S.C. Writing – Review & Editing: J.S-R., J.S-L., B.M., and S.K. Funding Acquisition: L.S.C. and S.K. Project administration: L.S.C. and S.K.

