## Supplementary figures and images for "SMAD4 and TGFβ are architects of inverse genetic programs during fate-determination of antiviral memory CD8 T cells"

### Supplementary Fig S1

Supplement Fig 1

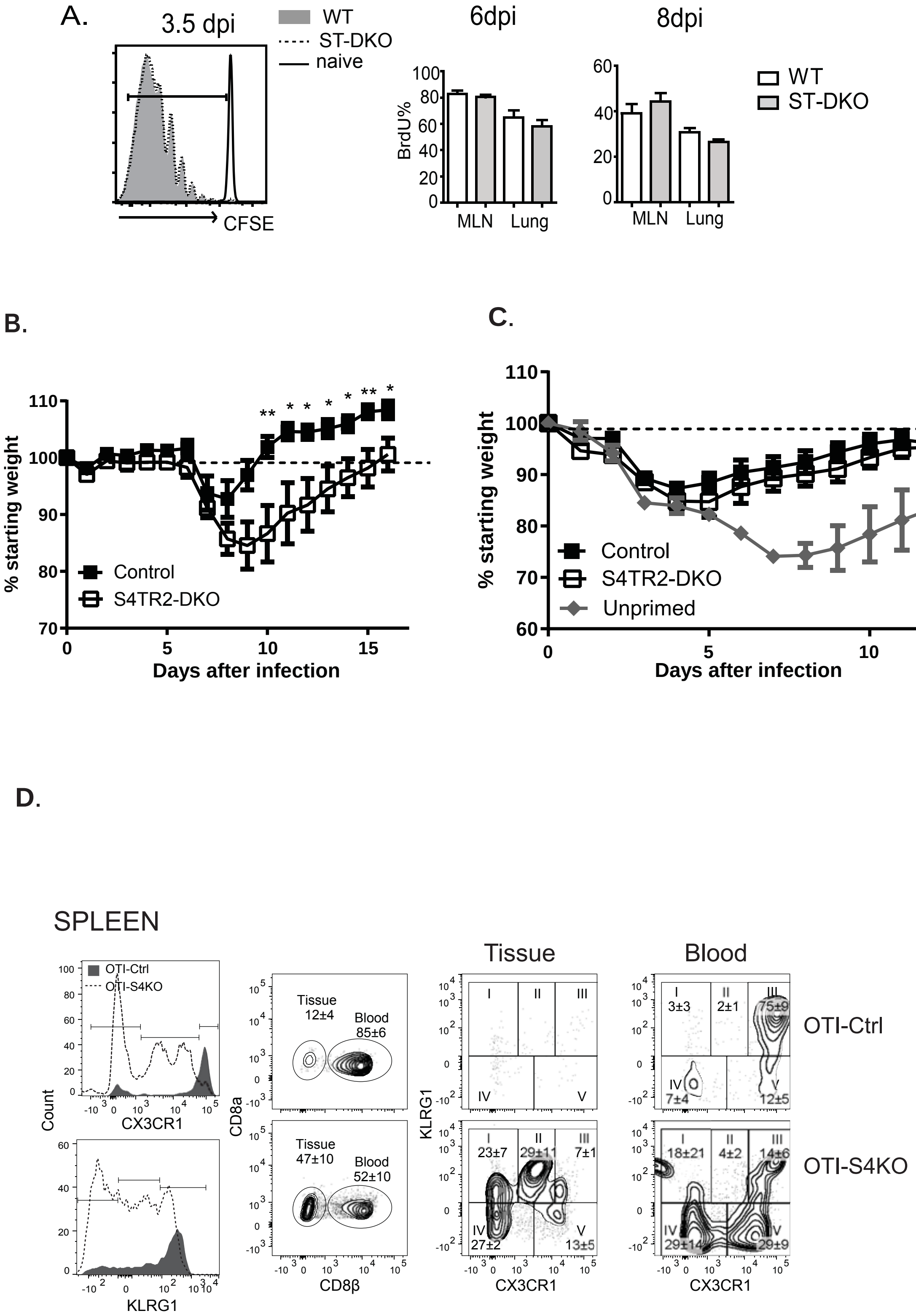

### Supplementary Fig S2

**Supplement Fig 2:**

**A.**

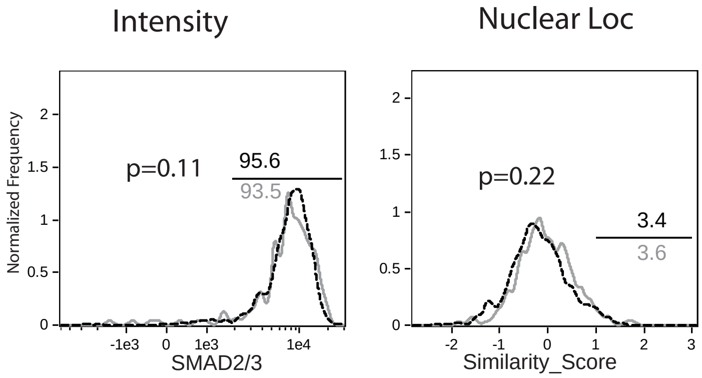

### Supplementary Fig S4

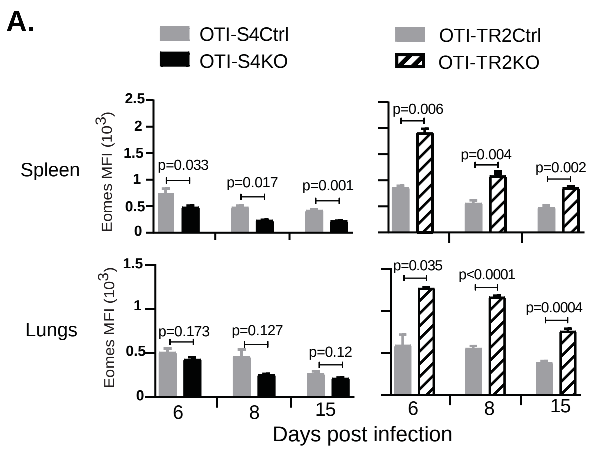

### Supplementary Fig S5

**Supplement Fig5:**

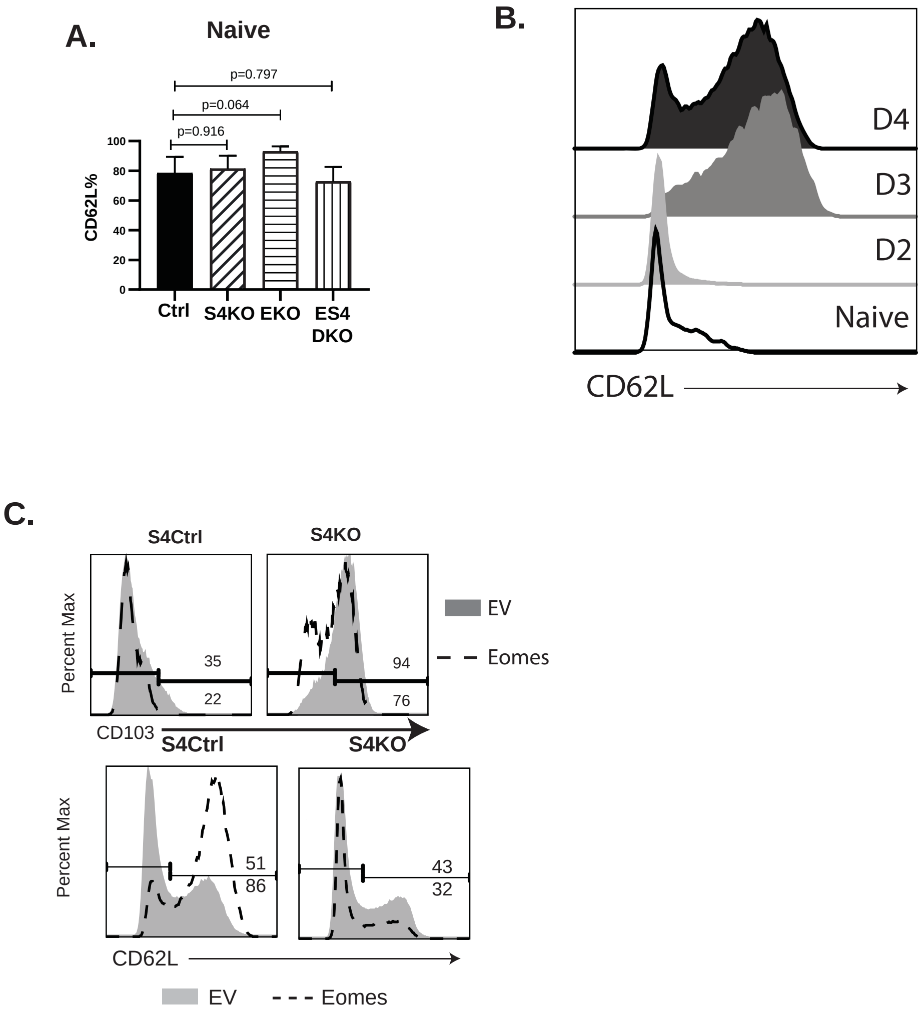
