## Supplemental Table for "SMAD4 and TGFβ are architects of inverse genetic programs during fate-determination of antiviral memory CD8 T cells"

Table1

| | | D6 | D8 | D10 | D15 | D30 |
|---|---|---|---|---|---|---|
| **KLRG1** | TKO | 0.0003 | <0.0001 | <0.0001 | <0.0001 | <0.0001 |
| | SKO | 0.2841 | <0.0001 | <0.0001 | <0.0001 | <0.0001 |
| | STDKO | 0.0517 | <0.0001 | <0.0001 | <0.0001 | <0.0001 |

| | | D6 | D8 | D10 | D15 | D30 |
|---|---|---|---|---|---|---|
| **CD127** | TKO | 0.0005 | <0.0001 | <0.0001 | <0.0001 | 0.0007 |
| | SKO | <0.0001 | <0.0001 | <0.0001 | <0.0001 | 0.0032 |
| | STDKO | <0.0001 | <0.0001 | <0.0001 | <0.0001 | 0.0012 |

| | | D6 | D8 | D10 | D15 | D30 |
|---|---|---|---|---|---|---|
| **CD62L** | TKO | <0.0001 | 0.8634 | 0.1047 | 0.0832 | 0.993 |
| | S4KO | 0.038 | 0.0033 | 0.2824 | 0.0003 | <0.0001 |
| | STDKO | 0.8727 | 0.0239 | 0.0499 | 0.0003 | <0.0001 |

| | | D6 | D8 | D10 | D15 | D30 |
|---|---|---|---|---|---|---|
| **CD103** | TKO | 0.0176 | <0.0001 | 0.8318 | 0.9376 | 0.0985 |
| | SKO | 0.0624 | <0.0001 | <0.0001 | <0.0001 | <0.0001 |
| | STDKO | <0.0001 | <0.0001 | <0.0001 | <0.0001 | <0.0001 |

Table2

| | | D5 | D7 | D14 |
|---|---|---|---|---|
| **KLRG1** | TR2KO | >0.9999 | 0.7471 | 0.041 |
| | S4KO | 0.008 | <0.0001 | <0.0001 |
| | TR2S4KO | 0.0016 | <0.0001 | <0.0001 |

| | | D5 | D7 | D14 |
|---|---|---|---|---|
| **CD127** | TR2KO | 0.021 | 0.0002 | 0.0025 |
| | S4KO | <0.0001 | <0.0001 | <0.0001 |
| | TR2S4KO | <0.0001 | <0.0001 | <0.0001 |

| | | D5 | D7 | D14 |
|---|---|---|---|---|
| **CD62L** | TR2KO | 0.0191 | 0.0406 | 0.0091 |
| | S4KO | <0.0001 | 0.0008 | <0.0001 |
| | TR2S4KO | <0.0001 | 0.0024 | <0.0001 |

| | | D5 | D7 | D14 |
|---|---|---|---|---|
| **CD103** | TR2KO | 0.0125 | 0.0006 | 0.002 |
| | S4KO | 0.0088 | <0.0001 | <0.0001 |
| | TR2S4KO | 0.2796 | <0.0001 | <0.0001 |
