## Supplementary legend for "SMAD4 and TGFβ are architects of inverse genetic programs during fate-determination of antiviral memory CD8 T cells"

**Supplemental Data.**

**Supplementary Figure S1.**

A) OTI-S4TR2Ctrl (grey shading) and OTI-S4TR2KO cells (dotted line) were labeled with CFSE and transferred to B6 mice before infection with X31-OVA. Histograms show CFSE-dilution at 3.5dpi.

OTI-S4TR2Ctrl (white fill) and OTI-S4TR2KO cells (grey fill) were transferred to B6 mice before infection with X31-OVA. Mice received BrdU by IP injection 3 hrs before sacrifice.

B) Mice were infected with X31-OVA (H3N2 serotype) and weighed daily. At 30dpi, the mice challenged with WSN-OVA_I_ (H1N1 serotype). Graphs show weight loss as % change from baseline (day 0). Symbols indicate statistical comparisons with controls P<0.05 (*), P<0.01 (**). Combined data from two independent experiments are shown. Weight change in S4TR2-Ctrl (filled squares) and S4TR2-DKO mice (open squares) after primary infection with X31-OVA

C) Weight change in S4TR2-Ctrl (filled squares) and S4TR2-DKO mice (open squares) after secondary infection with WSN-OVA_I_. UP are unprimed control mice.

D) B6 mice received mixed donor cells (OTI-S4KO and OTI-S4Ctrl) before infection with X31-OVA. Antibodies (CD8β) were injected 5 mins before sacrifice. Donor cells in spleens were analyzed for KLRG1 and GFP (CX_3_CR1) expression at 34dpi. Histograms donor cells from OTI-S4KO (dashed line) and OTI-S4Ctrl mice (continuous line). Contour plots show donor cells in the red (blood) and white (Tissue) pulp.

**Supplementary Fig. S2**

OTI-S4Ctrl and OTI-S4KO were transferred to C57BL/6 mice before infection with LM-OVA. Donor cells were analyzed for R-SMAD2/3 expression at 8 dpi using imaging flow cytometry. Similarity scores were determined using IDEAS software. Histograms show similar R-SMAD2/3 intensity and nuclear localization for OTI-S4Ctrl (grey line) and OTI-S4KO (dashed line) cells. Data are means ± SD, n=3.

**Supplementary Fig S4.**

Naïve CD8 T cells were transferred to B6 mice before infection with X31-OVA. On the days shown, CD45.1^+^ cells in the lungs and spleens were analyzed for EOMES expression by flow cytometry. Graphs show Eomes expression (MFI) in OTI-S4KO (black fill) and OTI-TR2KO (Cross hatch) cells, with the respective Cre-deficient controls (grey fill). Data are mean ± SD (n=4/group). Two independent experiments gave similar results.

**Supplementary Fig S5.**

A) Naïve CD8 T cells from S4KO, EKO, ES4KO and control mice express CD62L at similar levels

B) Naïve CD8 T cells were purified from SLO in C57BL/6 mice and stimulated with anti-CD3/CD28 and rIL-2 (48hrs) then washed and cultured with rIL-2 (48hrs) without TcR stimulation. Aliquots of CD8 T cells were analyzed for CD62L expression at the times indicated.

C) Overlaid histograms show CD103 (top) and CD62L (bottom) expression on S4KO and S4Ctrl cells after transduction with EVIRES-GFP (grey shading) EOMES-IRES-GFP (dashed lines)
